# Tick-borne pathogen detection in midgut and salivary glands of adult *Ixodes ricinus*

**DOI:** 10.1101/489328

**Authors:** Lejal Emilie, Moutailler Sara, Šimo Ladislav, Vayssier-Taussat Muriel, Pollet Thomas

## Abstract

**Background:** Tick midgut and salivary glands represent the primary organs for pathogen acquisition and transmission, respectively. Specifically, the midgut is the first organ to have contact with pathogens during the blood meal uptake, while salivary glands along with their secretions play a crucial role in pathogen transmission to the host. Currently there is little data about pathogen composition and prevalences in *I. ricinus* midgut and salivary glands. The present study investigated the presence of 32 pathogen species in the midgut and salivary glands of unfed *I. ricinus* males and females using high-throughput microfluidic real-time PCR. Such an approach is important for enriching the knowledge about pathogen distribution in distinct tick organs which should lead to a better understanding *I. ricinus*-borne disease epidemiology.

**Results:** *Borrelia lusitaniae, Borrelia spielmanii*, and *Borrelia garinii*, were detected in both midgut and salivary glands suggesting that the migration of these pathogens between these two organs might not be triggered by the blood meal. In contrast, *Borrelia afzelii* was detected only in the tick midgut. *Anaplasma phagocytophilum* and *Rickettsia helvetica* were the most frequently detected in ticks and were found in both males and females in the midgut and salivary glands. In contrast, *Rickettsia felis* was only detected in salivary glands. Finally *Borrelia miyamotoi* and *Babesia venatorum* were detected only in males in both midgut and salivary glands. Among all collected ticks, between 10 and 21% of organs were co-infected. The most common bacterial co-infections in male and female midgut and salivary glands was *R. helvetica/A. phagocytophilum* and *R. helvetica/B. lusitaniae* respectively.

**Conclusions:** Analysing tick-borne pathogen (TBP) presence in specific tick organs enabled us to (i) highlight contrasting results with well-established transmission mechanism postulates, (ii) venture new hypotheses concerning pathogen location and migration from midgut to salivary glands, and (iii) suggest other potential associations between pathogens not previously detected at the scale of the whole tick. This work highlights the importance of considering all tick scales (i.e. whole ticks *vs* organs) to study TBP ecology and represents another step towards improved understanding of TBP transmission.

## Introduction

Ticks are vectors of a large number of pathogenic microorganisms including bacteria, protozoa, and viruses, which cause serious diseases in both humans and animals. Ixodes ricinus is the predominant tick species in Europe and is recognised as the primary vector of infectious diseases in humans, including Lyme borreliosis caused by *Borrelia burgdorferi sensu lato*. Over the past decade, the number of studies characterizing pathogens carried by *I. ricinus* has considerably increased, enriching the wealth of information on *I. ricinus* pathogen composition (i.e. Reis *et al*., 2011, Estrada-Peña *et al*., 2011, Coipan *et al*., 2013, Reye *et al*., 2013, Jahfari *et al*., 2016, Moutailler *et al*., 2016, Marchant *et al*., 2017, Raileanu *et al*., 2017, Schotta *et al*., 2017, Szekeres *et al*., 2017, Chvostac *et al*., 2018). It’s also important to note that *I. ricinus* is regularly found to be co-infected by several pathogenic agents (Halos *et al*., 2005; Andersson *et al*., 2013; Berggoetz *et al*., 2014; Moutailler *et al*., 2016; Raileanu *et al*., 2017). This information is crucial considering that different co-infection combinations in humans and animals are highly likely to alter disease symptoms and severity (i.e. Grunwaldt *et al*., 1983; Golightly *et al*., 1989, Swanson *et al*., 2007, Diuk-Wasser *et al*., 2016). The vast majority of reports focusing on pathogen detection in *I. ricinus* have investigated intact bodies of adult ticks or pools of nymphs, however such methods cannot take into account organ-specific pathogen distribution. This information becomes significant as tick midgut and salivary glands act as specific and distinct barriers to efficient pathogen transmission, and are thus the main determinants influencing the acquisition, maintenance, and transmission of pathogens by ticks. Salivary glands may be co-infected by several different pathogens which can then be transmitted to the vertebrate host during blood feeding along with salivary secretions (Santos *et al*., 2002, Futse *et al*., 2003, Popov *et al*., 2007). Moreover, the tick midgut presents a pivotal microbial entry point and determines pathogen colonization and survival in the tick (Narasimhan *et al*., 2014). The common pathogenic lifecycle within the tick vector starts with ingestion of the infected host blood, pathogen migration through the gut to the haemocoel, transport into and infection of the salivary glands, then transmission to another host during the next blood meal. This pathway can vary depending on the pathogenic agent. For example, *Borrelia* and *Bartonella* species are known to be “stored” in the gut and migrate to the salivary glands during the next blood meal (see in Piesman and Schneider 2002), thus suggesting that blood meals trigger this migration from the midgut. In contrast, *Anaplasma* and *Ehrlichia* species are able to replicate in the midgut and migrate to the salivary glands in unfed ticks (Ueti *et al*., 2007). The infection/transmission cycle of *Babesia* species is similar to that of this second group, except that tick tissues are infected by different parasitic developmental stages (Hajdusek *et al*., 2013). While multiple different pathogens are now known to co-infect whole *I. ricinus* ticks, it is critical to identify pathogen presence at a finer scale in tick organs, to deepen our knowledge on pathogen associations and transmission and offer new insights into TBP and tick-borne diseases. By testing for the presence of 32 TBP species in both midgut and salivary glands of *I. ricinus* males and females using the microfluidic real-time PCR approach, we aim to detect pathogens in both of these key tick organs intimately involved in pathogen acquisition and transmission. Our study has consequently generated new hypotheses to understand TBP transmission.

## Material and Methods

### Tick collection and organ dissection

A total of 30 female and 30 male questing *I. ricinus* ticks were collected in May 2017 by flagging in the Senart forest (48°40’N, 2°29’E), south of Paris, France. Females and males were placed in separate sampling tubes. Before dissection, all ticks were washed once in 70% ethanol for 5 min and twice in distilled water for 5 min (Michelet *et al*., 2014). Tick organs, midgut and salivary glands, were then dissected in ice-cold PBS (pH = 7.2). The 120 samples were then conserved at -80°C until the DNA extraction.

### DNA extraction

Tissue samples were individually crushed with glass beads using the homogeniser Precellys®24 Dual at 5500 rpm for 20 s. Genomic DNA was then extracted using the Nucleospin tissue DNA extraction Kit (Macherey-Nagel, France). For each sample, total DNA was eluted in 50 μL of rehydration solution and stored at -20°C until further analysis.

### High-throughput screening of bacterial and parasitic tick-borne pathogens

The BioMark^™^ real-time PCR system (Fluidigm, USA) was used for high-throughput microfluidic real-time PCR for the most common bacterial and parasitic TBP species known to circulate or emerge in Europe. For bacteria, seven species belonging to the Lyme spirochete group (*Borrelia burgdorferi sensu lato*) were tested: *B. burgdorferi sensu stricto, B. afzelii, B. garinii, B. spielmanii, B. valaisiana, B. lusitaniae*, and *B. bissettii*. We also tested one species belonging to the *Borrelia* recurrent fever group, *B. miyamotoi*. Six species of *Anaplasma: A. phagocytophilum, A. platys, A. marginale, A. ovis, A. centrale*, and *A. bovis*. Six species of *Rickettsia* from the spotted fever group: *R. felis, R. helvetica, R. conorii, R. slovaca, R. massiliae*, and *R. aeschlimannii. Ehrlichia canis, Candidatus* Neoehrlichia mikurensis, *Bartonella henselae, Francisella tularensis, Coxiella burnetii*, and *Mycoplasma spp*. were also tested. For parasites (Apicomplexa), seven species of *Babesia* were tested: *B. canis, B. ovis, B. microti, B. bovis, B. caballi, B. venatorum*, and *B. divergens*. We also tested *Theileria* spp, and *Hepatozoon* spp. Extraction and amplification controls were included using DNA extracts from *I. ricinus* and *E. coli* respectively.

As previously described in Moutailler *et al* (2016), a DNA pre-amplification step was performed for each sample in a final volume of 5 μL containing 2.5 μL TaqMan PreAmp Master Mix (2X), 1.2 μL of the pooled primer mix (0.2X), and 1.3 μL of tick DNA, with one cycle at 95°C for 10 min, 14 cycles at 95°C for 15 s, followed by 4 min at 60°C. Following pre-amplification, qPCRs were performed using FAM- and black hole quencher (BHQ1)- labelled TaqMan probes (Michelet *et al*., 2014) with TaqMan Gene Expression Master Mix in accordance with manufacturer’s instructions (Applied Biosystems, France). Thermal cycling conditions were as follows: 95°C for 5 min, 45 cycles at 95°C for 10 s, 60°C for 15 s, and 40°C for 10 s. Data were acquired on the BioMark^™^ Real-Time PCR system and analysed using the Fluidigm Real-Time PCR Analysis software to obtain crossing point (CP) values.

## Results

Of the collected ticks, 73% of males and 59% of females were infected by at least one pathogen (Figure 1), and of all *Anaplasma* and *Rickettsia* species tested in the study, only *A. phagocytophilum, Rickettsia helvetica*, and *Rickettsia felis* were detected (Figure 2, Table 1). *A. phagocytophilum* and *R. helvetica* were detected in both salivary glands and midguts in males and females. *A. phagocytophilum* was detected in 23% of male organs, and 17% of female midgut and salivary glands (Figure 2). *R. helvetica* infection rates reached 27% and 20% in male midgut and salivary glands respectively, whereas in females, infection rates were 14% in both midgut and salivary glands. Both *A. phagocytophilum* and *R. helvetica* were mostly simultaneously detected in both organs, and were occasionally found in either only the midgut or salivary glands (Table 1). *Rickettsia felis* was detected only in salivary glands in males and females with an infection rate reaching 7% in this organ. Members of the complex *B. burgdorferi s.l*. (*B. lusitaniae, B. spielmanii, B. afzelii*, and *B. garinii*) were detected only in females. *B. lusitaniae, B. spielmanii*, and *B. garinii* were detected in both midgut (7%, 7%, and 3% respectively) and salivary glands (7%, 3%, and 3% respectively) (Figure 2, Table 1). *B. afzelii* was only detected in 7% of female midguts. Note that the other members of the complex tested in this study, *B. burgdorferi s.s., B. valaisiana*, and *B. bissettii* were not detected in either males or females. *B. miyamotoi* was detected only in males, and mostly in salivary glands (13%), rarely in both organs (3%) and never in the midgut alone (Figure 2, Table 1). The other bacterial species tested in this study, *Ehrlichia canis, Candidatus Neoehrlichia mikurensis, Bartonella henselae, Francisella tularensis, Coxiella burnetii* and *Mycoplasma* spp were not detected in our samples. Apicomplexa were detected in both males and females in both MG and SG. Among all tested parasites, only *Babesia venatorum* was only detected in males (Figure 2, Table 1) with 7% infected in both organs, while 13% were infected only in salivary glands.

**Figure 1:**
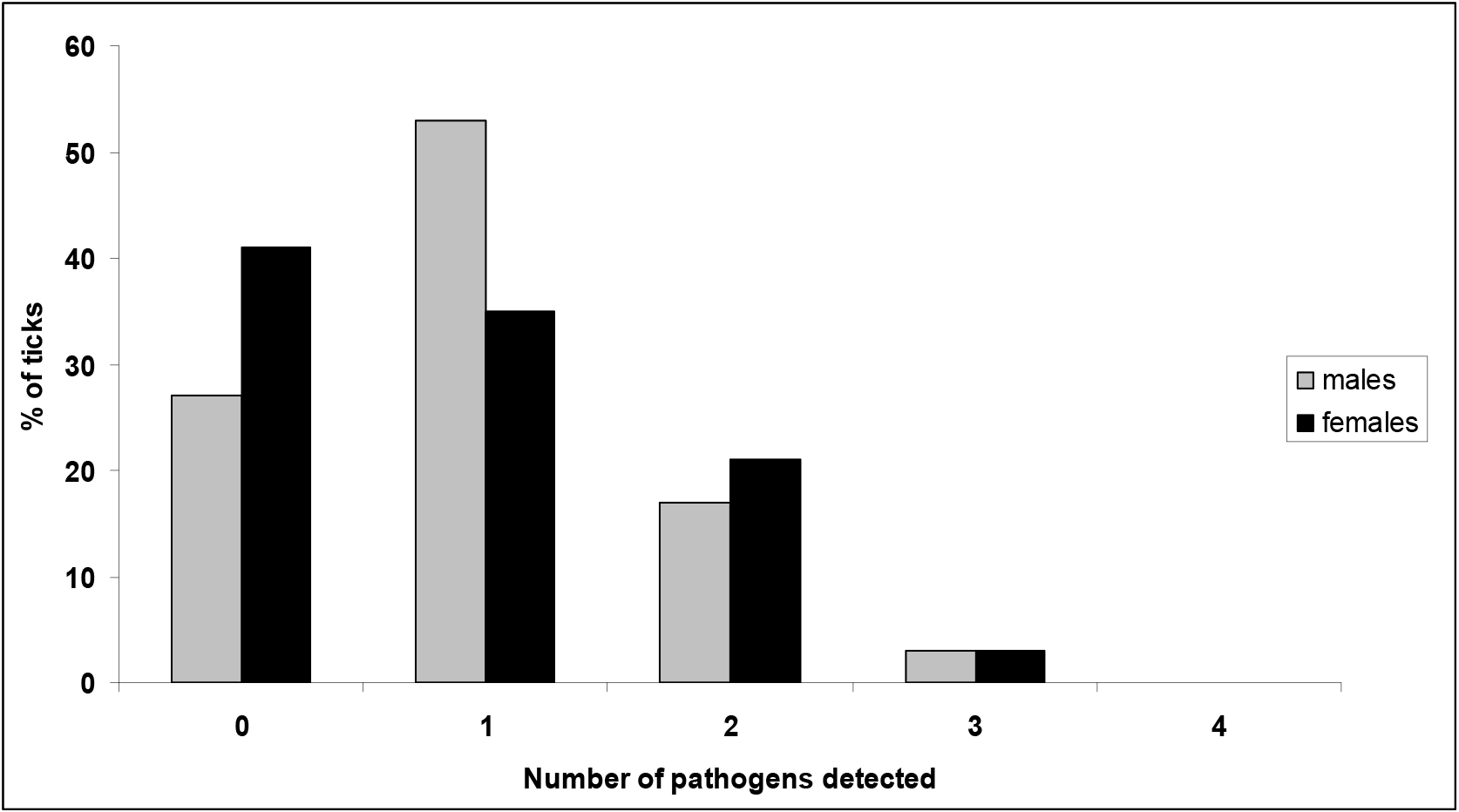
Pathogen infection rates (%) in *Ixodes ricinus* males and females, according to the number of pathogens detected.

**Figure 2:**
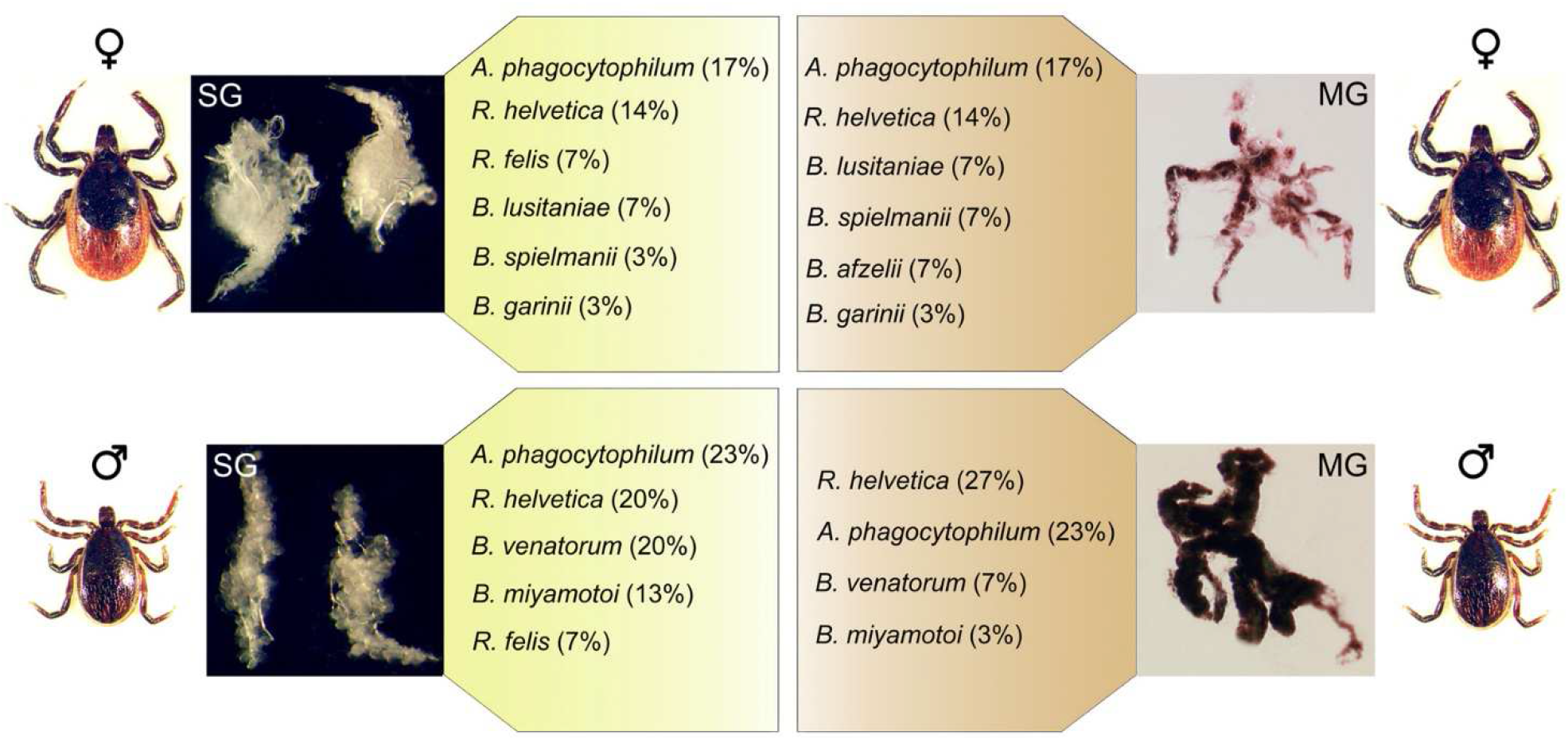
Pathogen infection rates (%) in salivary glands (SG) and midgut (MG) of unfed *I. ricinus* females (♀) and males (♂).

**Table 1:** Pathogen infection rates (%) in females (F) and males (M) either in salivary glands alone (SG), in midgut alone (MG), or in both organs (SG/MG). ND = Not Detected.

|  | F |  |  | M |  |  |
| --- | --- | --- | --- | --- | --- | --- |
|  | SG | MG | SG/MG | SG | MG | SG/MG |
| *A. phagocytophilum* | 7 | 7 | 10 | 3 | 3 | 20 |
| *R. helvetica* | 3 | 3 | 10 | 7 | 13 | 13 |
| *R. felis* | 7 | ND | ND | 7 | ND | ND |
| *B. lusitaniae* | ND | ND | 7 | ND | ND | ND |
| *B. spielmanii* | ND | 3 | 3 | ND | ND | ND |
| *B. afzelii* | ND | 7 | ND | ND | ND | ND |
| *B. garinii* | ND | ND | 3 | ND | ND | ND |
| *B. miyamotoi* | ND | ND | ND | 10 | ND | 3 |
| *B. venatorum* | ND | ND | ND | 13 | ND | 7 |

It is also important to note that 20% of males and 24% of females were co-infected. In males, 10% of midgut and 17% of salivary gland samples were co-infected (Figure 3). In females, 17% of midgut and 21% of salivary gland samples were co-infected (Figure 3). The most common co-infection in both salivary glands and midgut of males and females was *R. helvetica/A. phagocytophilum and R. helvetica/B. lusitaniae* respectively.

**Figure 3:**
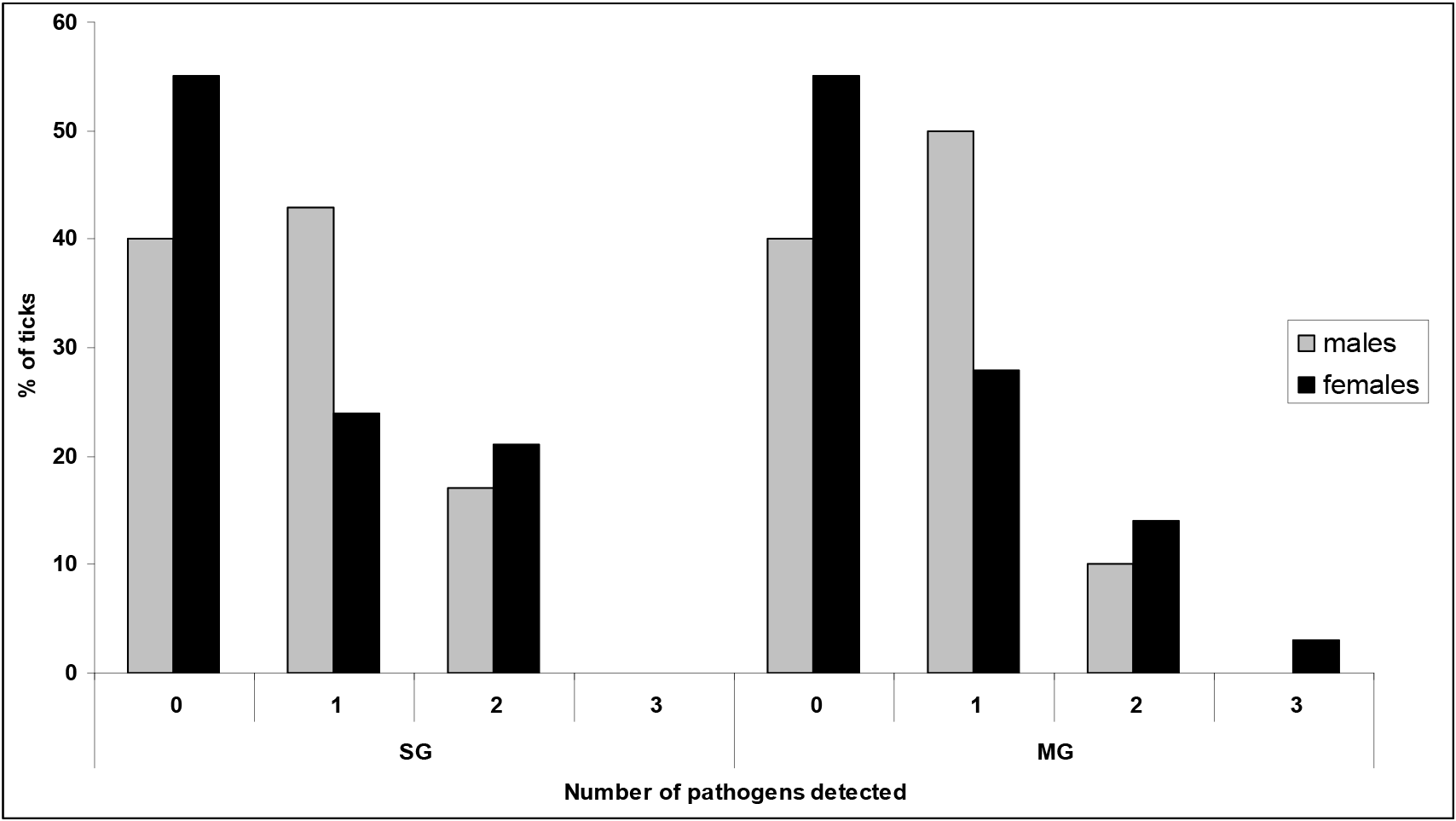
Pathogen infection rates (%) in midgut (MG) and salivary glands (SG) of unfed *I. ricinus* males and females according to the number of pathogens detected.

## Discussion

### Results contrasting with well-established postulates

Several *B. burgdorferi sensu lato* genospecies (*B. lusitaniae, B. spielmanii*, and *B. garinii*) were only detected in questing females. This result supports previous findings showing that female ticks in woodland sites were more likely to be infected by *B. burgdorferi sensu lato* than males and nymphs (Halos *et al*., 2010) suggesting potential interactions between female ticks and these bacteria. However the most unexpected results was that we detected these species in both salivary glands and midgut which contrasts with the well-established postulate that *B. burgdorferi sensu lato* genospecies are not found in salivary glands during the initial tick attachment, as they only move rapidly from the gut to the salivary glands at the beginning of the next blood meal (Des Vignes *et al*., 2001, Piesman *et al*., 2001; Kung *et al*., 2013). Our findings would suggest that some *B. burgdorferi sensu lato* genospecies do not need a blood meal to start their multiplication and migration from the gut to salivary glands. This hitherto unseen observation on field-collected ticks is completely consistent with the recent experimental results of Sertour *et al* (2018). These authors detected *Borrelia burgdorferi* sl strains in female salivary glands prior to blood meals, and also showed that these bacteria could infect mice within 24 h of tick bite. We also note that in our study, the presence of these different *B. burgdorferi sl* genospecies in salivary glands could also be explained if not all bacteria were transmitted to the host, and that residual bacteria thus remained in salivary glands after moulting. In this case, their persistence in the salivary glands of unfed ticks would suggest that this organ could also be, as for the midgut, a potential reservoir for these spirochetes. Whichever the hypothesis, this result contributes important information to the understanding of tick-borne pathogen transmission as it suggests that several species of the *Borrelia burgdorferi sl* complex could be rapidly transmitted after tick bite.

### Pathogen location and migration in ticks

Interestingly—and as has been previously observed for the other *B. burgdorferi sl* genospecies—*B. afzelii* were only detected in females, and exclusively in the midgut. These results are highly consistent with the study by Pospisilova *et al* (2018) in which *Borrelia afzelii* were abundant in the guts of unfed *I. ricinus* nymphs, whereas the spirochetes were not present in salivary glands. This result better corresponds to the previously-cited postulate and thus contrasts with the findings discussed for the other *B. burgdorferi sensu lato* genospecies. This result advocates greater caution when generalising transmission mechanisms, which may differ according to *B. burgdorferi sensu lato* genospecies.

We detected *R. helvetica* and *A. phagocytophilum* in males and females in both salivary glands and midgut, and both of these pathogenic agents were the most frequently detected in the collected ticks. Their ubiquity and high proportions in tick organs suggest that they could be well adapted to the different environmental conditions characterising both salivary glands and midgut, and thus may be stronger competitors than other pathogen species. Several studies have shown low *A. phagocytophilum* prevalence or titter in the salivary glands of unfed ticks (e.g. Alberdi *et al*., 1998) suggesting that pre-feeding ticks may induce bacterial replication. As “pre-feeding” is probably a rare event in the field, and that it is unlikely that all the questing ticks infected by *A. phagocytophylum*—approximately 20% of infected ticks— were pre-fed, our results suggest that this bacteria could already be present in the salivary glands of unfed ticks at high infection rates. On the other hand, detecting *R. helvetica* in questing *I. ricinus* is not surprising as their potential presence in this tick species is well established. However, no clear evidence of their transmission by ticks is currently available. Detecting this pathogen in salivary glands is an important epidemiological result as it represents an additional indirect argument that this pathogen could be transmitted by tick bites. For each tick infected by *A. phagocytophilum* and *R. helvetica*, these bacteria were most often simultaneously detected in both midgut and salivary glands and only sometimes in either the midgut or salivary glands alone. These results are important in terms of pathogen transmission as they suggest that not all bacterial cells migrate together from the midgut to salivary glands, and that some infected females (with pathogens only in the midgut) would not necessarily be infectious. In contrast to *R. helvetica*, we only detected *R. felis* in the salivary glands of both males and females. *R. felis* is known to be mainly transmitted from cat to cat *via* fleas, with human contamination arising from cat or flea bites. Already detected in engorged *Rhipicephalus sanguineus* (Oliveira *et al*., 2008; Abarca *et al*., 2013), two studies have also identified this pathogen in questing *I. ricinus* nymphs (Vayssier-Taussat *et al*., 2013, Lejal *et al*., unpublished). Detecting this pathogen only in salivary glands could suggest that it doesn’t remain in the midgut and rapidly migrates to the salivary glands. This result also implies that *R. felis* would already be present in salivary glands at the time of the next blood feeding and thus might be rapidly transmitted after tick bite. While the presence and prevalence of this pathogenic agent are rarely investigated in studies dealing with tick-borne pathogens, this new detection in *I. ricinus*, and specifically in the salivary glands, should encourage us to increase our surveillance for this pathogen causing spotted fever in humans. Future investigations should be carried out to clarify the potential vector competence of *I. ricinus* for this pathogen and its role in transmission.

*Borrelia miyamotoi* of the *Borrelia* recurrent fever group and the parasite *Babesia venatorum* were only detected in males. That they were detected in general was not surprising as they have already been identified in many European countries (i.e. Fraenkel *et al*., 2002; Richter *et al*., 2003; Geller *et al*., 2012; Reye *et al*., 2013; Cosson *et al*., 2014; Sakakibara *et al*., 2016; Moutailler *et al*., 2016; Paul *et al*., 2016; Oechslin *et al*., 2017; Szekeres *et al*., 2017; Wagemaker *et al*., 2017). Interestingly, these pathogenic agents were found in both salivary glands and midgut, suggesting—particularly for *B. miyamotoi*—that migration from midgut to salivary glands is not triggered by a blood meal. Detecting these pathogens in salivary glands is particularly significant in terms of public and animal health. Indeed, in the event that females are infected with these pathogens (i.e. Moutailler *et al*., 2016, Diaz *et al*., 2017), it seems that both organs could be potential reservoirs for these pathogenic agents, thus potentially facilitating rapid transmission after tick bite. It will of course be necessary to verify whether the *B. venatorum* parasites residing in salivary glands are actually infectious.

### Potential associations between pathogens

We note finally that among all infected ticks, co-infections were observed in 20% of males and 24% of females. At a finer scale, between 10 and 21% of organs were co-infected. With detection tools becoming more sensitive and efficient, tick co-infections are observed with greater frequency (Halos *et al*., 2005, Schicht *et al*., 2011, Andersson *et al*., 2013, Cosson *et al*., 2014, Castro *et al*., 2015, Moutailler *et al*., 2016, Raileanu *et al*., 2017), suggesting that tick co-infection is the rule rather than the exception (Moutailler *et al*., 2016). In this study, this hypothesis could also be extended to the level of individual tick organs. These observations—particularly co-infection in salivary glands—are particularly relevant to public health, as the tick disease community is starting to accept that pathogen co-transmission might significantly modify disease symptoms and severity (i.e. Krause *et al*., 1996, Swanson *et al*., 2007, Diuk-Wasser *et al*., 2016). The most common bacterial co-infection in male salivary glands and midgut was *R. helvetica/A. phagocytophilum*, whereas in females, the most frequent association in salivary glands and midgut was *R. helvetica/B. lusitaniae*. Our data contrast with previous findings performed in whole ticks demonstrating that the most common *I. ricinus* co-infection was *B. garinii* and *B. afzelii* (i.e. Moutailler *et al* 2016, Raileanu *et al* 2018). Analysing TBP co-infections at a finer scale clearly enables us to highlight other potential associations not detected at the level of the whole tick.

## Conclusions

Analysing tick-borne pathogen composition and infection rates at the scale of individual tick organs highlighted (i) some pathogen location results which contrasted with well-established transmission mechanism postulates, (ii) novel findings in terms of pathogen location and migration time from midgut to salivary glands, and (iii) other potential associations between pathogens not detected at the scale of whole ticks. All findings should be confirmed using RNA extracts in order to verify if pathogens are indeed viable and actively replicating, to validate our hypotheses. This study represents another step towards generating improved control methods for tick-borne diseases and highlights the importance of investigating TBP infection at all scales (i.e. whole ticks vs organs) to better understand the ecology and epidemiology of tick-borne diseases. To identify the microbiota influence on pathogens, it would be interesting to characterise the microbiota in both *I. ricinus* male and female salivary glands and midgut to detect potential co-occurrences between pathogens and other microbes.

## Competing interests

The authors declare that they have no competing interests

## Funding

French National Institute for Agricultural Research, Animal Health department (France).

## Author contributions

Conceived and designed the experiments: TP, MVT. Performed the experiments: EJ, LS, TP. Analysed the data: EJ, TP, SM. Wrote the paper: EJ, SM, LS, MVT, TP

## Acknowledgements

We thank the VECTOTIQ team members in the BIPAR unit for stimulating discussion and support.

